# Natural Variation in HIV-1 Entry Kinetics Map to Specific Residues and Reveal an Interdependence Between Attachment and Fusion

**DOI:** 10.1101/2024.06.25.600587

**Authors:** Nicholas E. Webb, Colin M. Sevareid, Carolina Sanchez, Nicole H. Tobin, Grace M. Aldrovandi

## Abstract

HIV-1 entry kinetics reflect the fluid motion of the HIV envelope glycoprotein through at least three major structural configurations that drive virus-cell membrane fusion. The lifetime of each state is an important component of potency for inhibitors that target them. We used the time-of-addition inhibitor assay and a novel analytical strategy to define the kinetics of pre-hairpin exposure (using T20) and co-receptor engagement (via. maraviroc), through a characteristic delay metric, across a variety of naturally occurring HIV Env isolates. Among 257 distinct HIV-1 envelope isolates we found a remarkable breadth of T20 and maraviroc delays ranging from as early as 30 seconds to as late as 60 minutes. The most extreme delays were observed among transmission-linked clade C isolates. We identified four single-residue determinants of late T20 and maraviroc delays that are associated with either receptor engagement or gp41 function. Comparison of these delays with T20 sensitivity suggest co-receptor engagement and fusogenic activity in gp41 act cooperatively but sequentially to drive entry. Our findings support current models of entry where co-receptor engagement drives gp41 eclipse and have strong implications for the design of entry inhibitors and antibodies that target transient entry states.

**Author Summary.:** The first step of HIV-1 infection is entry, where virus-cell membrane fusion is driven by the HIV-1 envelope glycoprotein through a series of conformational changes. Some of the most broadly active entry inhibitors work by binding conformations that exist only transiently during entry. The lifetimes of these states and the kinetics of entry are important elements of inhibitor activity for which little is known. We demonstrate a remarkable range of kinetics among 257 diverse HIV-1 isolates and find that this phenotype is highly flexible, with multiple single-residue determinants. Examination of the kinetics of two conformational landmarks shed light on novel kinetic features that offer new details about the role of co-receptor engagement and provide a framework to explain entry inhibitor synergy.

## Introduction

HIV-1 entry is driven by the trimeric envelope glycoprotein (Env) through a series of coordinated structural rearrangements in its two subunits, gp120 and gp41 (Fig. 1a). Transitional conformations that emerge during this process are the targets of broadly active fusion and anchor inhibitors. Because these targets are transient, the kinetics of entry and the lifetime of their exposure are important components of inhibitor activity and resistance[1–5].

**Figure 1:**
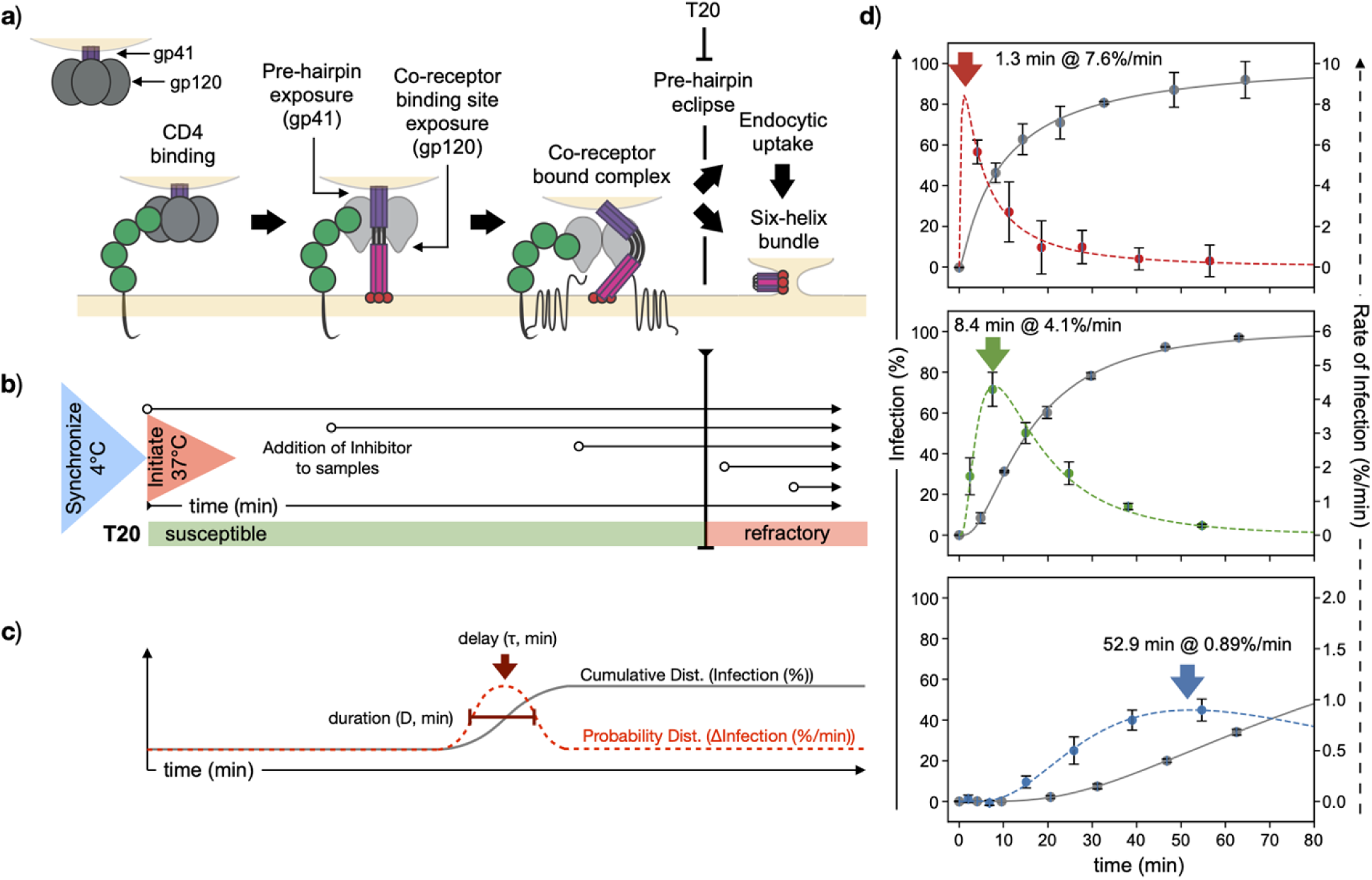
Time of Addition Kinetics Assay and Lognormal Fitting. (a) Model of HIV-1 entry depicting the conformational states of gp120 and gp41, and the pre-hairpin target of T20. (b) Time of addition schematic where entry is synchronized and initiated with a spinoculation and temperature shift. At various times, a saturating dose of inhibitor is added (arrows). Entry is susceptible until it progresses to the inhibitor-refractory state, which corresponds to the inhibitor’s target. (c) Cumulative distribution of refractory transitions (grey solid line, % infection) and time-dependent probability distribution (red dashed lines, Δinfection). The probability distribution is characterized by the timing of its peak rate (delay, τ) and width (duration, D). (d) Examples of early (top), moderate (middle) and late (bottom) T20-refractory transitions with delays and peak rates indicated. Cumulative infection (grey circles) and rate of infection (red, green and blue circles) are shown with corresponding lognormal fits. Data points and error bars in (d) indicate the average and standard deviation of duplicate measurements from a single representative experiment.

Entry is initiated when gp120 binds to its primary receptor CD4. This induces conformational changes that allow gp120 to bind one of two chemokine co-receptors, CCR5 or CXCR4[6]. CD4-binding also triggers the exposure of gp41[7–11], where the fusion peptides of gp41 are cast outward and become embedded in the cell membrane[1, 12]. In this elongated, pre-hairpin state, N-terminal and C-terminal heptad repeat regions of gp41 (NHR and CHR) are exposed and then fold together into a six-helix bundle that induces virus-cell fusion (Fig. 1a). During collapse or exposure of the pre-hairpin HIV-1 may also undergo endocytosis, blocking access of the pre-hairpin structure and downstream intermediates from inhibitors[13, 14].

The fusion peptide, NHR and CHR are all targets of broadly active inhibitors and antibodies[1, 15–17]. T20, the first clinically approved fusion inhibitor, targets the NHR and blocks six-helix bundle formation[18]. Because the NHR is only exposed during the pre-hairpin state (Fig. 1a), the sensitivity of an Env to T20 is partially determined by the lifetime of pre-hairpin exposure[19–21]. Thus, the overall kinetics of entry and the lifetime of key intermediates are important elements of fusion inhibitor potency and resistance[1].

Kinetics can also be linked to synergy between co-receptor antagonists and fusion inhibitors. For example, low CCR5 density and co-receptor antagonists slow entry kinetics [19, 20] and increase the potency of fusion inhibitors[22–25]. Thus, co-receptor engagement may resolve CD4-induced pre-hairpin exposure either by triggering six-helix bundle formation or endocytic uptake[13, 14]. Delaying this process, conversely, increases the duration of pre-hairpin exposure and its sensitivity to T20. Resistance to fusion inhibitors can also involve off-target substitutions associated with CD4/CCR5 engagement[2–5]. Thus, co-receptor engagement may serve as an alternate pathway of resistance.

Our current knowledge of HIV entry kinetics is largely restricted to a small number of HIV-1 Env isolates[19–21, 26–29]. HIV-1, however, is a highly mutable and structurally labile virus that can develop unique, closely related phylogenetic lineages within a single individual[30, 31]. In this study we wanted to understand the breadth of HIV-1 entry kinetics that exist naturally, among a large and diverse panel of isolates. We also wanted to evaluate the potential role of kinetics in transmission and acute viral adaptation and to examine mechanistic relationships between co-receptor engagement and pre-hairpin exposure in a broad context. To do this, we employed a panel of 257 diverse HIV-1 Env isolates with broad geographic, phylogenetic and clinical characteristics. These include a globally representative panel of tier 2 isolates[32], acutely transmitted clade B isolates [33] and isolates associated with adult and mother-to-child transmission[31, 34–39].

To accommodate high-volume kinetic profiling we streamlined the time of addition inhibitor assay, which measures the emergence of inhibitor-refractory virions as they progress through entry. We established a distributive model able to recapitulate entry kinetics data through characteristic delay (𝜏) and duration (D) metrics, where the delay marks the time of the inhibitor-refractory transition. Using this analytical strategy we identified a remarkable breadth of delays for T20 and the co-receptor antagonist maraviroc (MVC) ranging from as early as 30 seconds to as late as 60 minutes. Further, we identified single-residue determinants of entry kinetics that are associated with either attachment or fusion. Pairwise comparison of MVC and T20 delays reveal a correlation between kinetics and T20 sensitivity that suggests co-receptor engagement and the pre-hairpin drive fusion and/or endocytosis in a mutual fashion. This interplay suggests compensatory mechanisms in the adaptive pathways associated with fusogenicity and co-receptor engagement, which represent a new dimension to co-receptor usage efficiency that will be important for understanding resistance to fusion and anchoring inhibitors.

## Results

### The Lognormal Distribution Captures the Timing of Entry Transitions

The time of addition inhibitor assay measures a population of synchronized virions as they transition from an inhibitor-susceptible state to an inhibitor-refractory state (Fig. 1). Entry is synchronized and initiated by 4°C spinoculation followed by a rapid 37°C temperature shift. Susceptibility is tested over time by inhibitor saturation (Fig. 1b), where the inhibitor used determines which state of entry is being assessed. For example, entry is susceptible to T20 until its target, the pre-hairpin, is no longer accessible due to either its collapse into the six-helix bundle or endocytic uptake [13] (Fig. 1b). Thus, increases in T20-refractory infection over time reflect a general eclipse of the pre-hairpin state.

The time to 50% refractory infection (t_50_) is a standard timing metric based on the synchronized nature of the refractory transition. As a more empirical metric that does not attempt to model the entry process in detail, it can be used to measure discrete entry states independently without requiring additional rate parameters. The t_50_ is based on a model in which the synchronized refractory transitions follow a normal probability distribution (Fig. 1c, red dotted line). The peak of this distribution is the time, post initiation, that the largest fraction of virions transition, which we call the delay (𝜏, Fig. 1c). This distribution also has a width, or duration (D, full width in time at 50% peak rate, Fig. 1c) that describes the synchronicity with which the transition occurred. Because each time point of the time-of-addition assay is a cumulative measure of refractory infection (Fig. 1c, grey solid line), the corresponding probability distribution can be directly evaluated by differentiating raw refractory infection data over time.

Fig. 1d (grey circles) shows examples of early (top), moderate (middle) or late (bottom) T20-refractory transitions using the time-of-addition assay. We assumed that these data represented cumulative distributions and calculated the corresponding probability distributions directly by differentiation (Fig. 1d, red, green and blue circles). These data formed right-tailed probability distributions (dashed lines) consistent with a lognormal distribution function. Accordingly, we found the lognormal distribution to be a highly representative fit of the raw time-of-addition results, with R^2^>0.9 for 94% of all 712 kinetic data sets measured for this study (Fig. S1a).

Peak infection rates generally occurred before 50% cumulative infection (Fig. 1d), thus, t_50_ correlated poorly with the lognormal delay (Fig. S1d) while the time to 20% infection (t_20_) was more reflective (Fig. S1e). Both t_50_ and t_20_ are confounded by incomplete inhibition [40–42] and infection that occurs before initiation. This baseline infectivity does not, however, affect the differential delay metric. Overall baselines, however, were within ±3% infection for 93% of all data sets (Fig. S1f and Fig. S1g). Interestingly. while T20 baselines tended to be negative, MVC baselines tended to be positive.

We find that the lognormal distribution accurately represents the inhibitor-refractory transition through delay (𝜏) and duration (D) metrics and offers additional details regarding transitional synchronicity. Like t_50_, the lognormal distribution makes no assumptions about the mechanism of entry and does not require kinetic characterization of prerequisite entry stages. Unlike t_50_, the lognormal distribution can be used to obtain a high-quality fit of time-of-addition curves.

### Natural Breadth of Entry Kinetics

HIV entry is a concerted succession of states where later steps are delayed by lags in early steps. we used the T20 delay, which measures one of the more terminally accessible states of entry, as a general measurement of overall entry kinetics. To delineate the natural breadth of kinetics we measured T20 delays for 257 primary HIV-1 Env isolates including a globally representative panel of tier 2 isolates[32], acute clade B transmitter/founder isolates [33] and isolates associated with adult [34–36] and mother-to-child [31, 37–39] transmission.

Among all 257 isolates, we observed a broad distribution of T20 delays ranging from as little as 30 seconds to as late as 60 minutes (Fig. 2a). Due to physical limitations, delays <2 min were inconsistent across experiments, having coefficients of variation as high as 140%. Given this, we defined 2 minutes as our limit of quantitation. Delays of 2-4 min had 90% confidence intervals as high as 2.5-fold of the mean delay and all delays later than 4 minutes had 90% confidence intervals below 2-fold of their mean (Fig. S1f). Based on this two-fold confidence interval, we categorized T20 delays as extreme early (<2 min, Fig. S2a), early (2-4 min, Fig. S2b), typical (4-16 min), late (16-32 min, Fig. S3a) or extreme late (>32 min, Fig. S4b). Overall, 81% were typical and were evenly divided between 4-8 and 8-16 min. Less than 1% were extreme early and 10% were early. Roughly 6% were late and 2% were extreme late (>32 min). All delays are described in Table S1.

**Figure 2:**
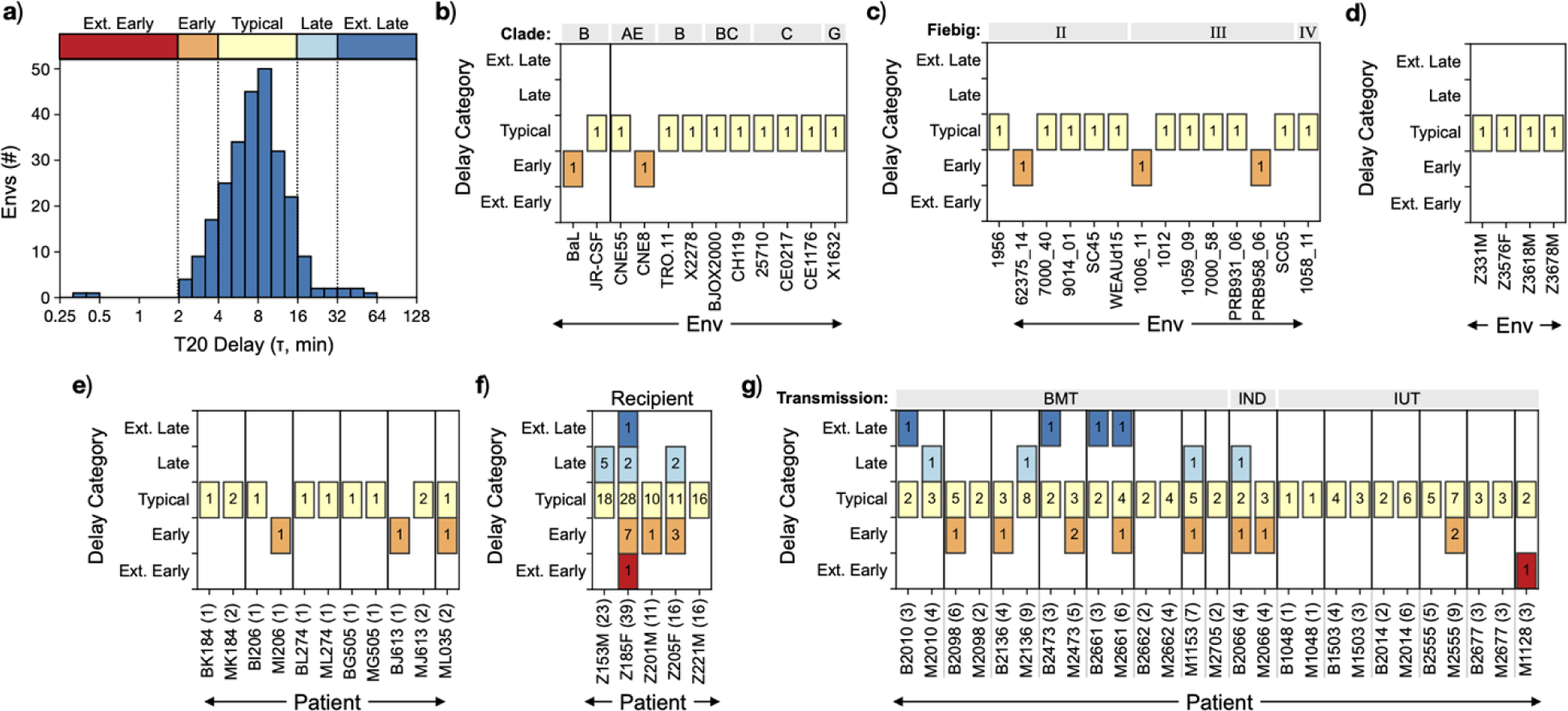
Broad Diversity of T20 Delay. (a) Distribution of T20 delays among 257 distinct HIV-1 isolates categorized into extreme early (<2 min), early (2-4 min), typical (4-16 min), late (16-32 min) and extreme late (≥32 min) ranges. (b-d) T20 delay categories for (b) a globally representative panel of tier 2 isolates (clade indicated above graph area, BaL and JR-CSF are added to this panel), (c) acute transmitter/founder isolates (clade B, Fiebig stage indicated above graph area) and (d) clade C transmitter/founder consensus isolates. Each Env in (b-d) was a single Env isolated from a single individual. (e) T20 delay categories for vertically transmitted isolates that include one infant isolate and up to two maternal isolates from each mother/infant pair (vertical lines). All isolates in (e) are clade A except for BK184/MK184, which are clade C/D and ML035, which is clade D/A. (f-g) T20 delay categories for (f) clade C isolates sampled longitudinally from adults within the first year of infection and (g) paired infant and maternal clade C isolates associated with vertical transmission. Multiple Envs were isolated from individuals in (e-g). Numbers within each box indicate the number of isolates, from each individual, belonging to the respective T20 delay category. Numbers in parentheses next to id’s indicate the number of Envs tested from each subject. Breast milk (BMT), in utero (IUT) and indeterminate (IND) transmission routes are indicated in (g) (above graph area). All T20 delays are the average of 1-7 independent experiments, each consisting of duplicate measurements, with lognormal R^2^≥0.9.

The T20 delay for HIV-1 BaL (clone 26) fell on the edge of the early/typical range (3.9 ± 2.17 min, SD, n=7) and was similar to the typical delay of JR-CSF (4.5 ± 1.49 min, SD, n=3, Fig. 2b). Among isolates from the global neutralization panel [32] (Fig. 2b), only CNE8 had an, early delay of 2.7 ± 0.21 min (SD, n=4). CNE8 is considered a highly sensitive tier 2 Env that exhibits a tier 1 phenotype when assessed using different cell lines[32], suggesting early delay may be associated with neutralization sensitivity. X2278, however, is also considered highly sensitive for the same reason and had a much later T20 delay of 9.9 min (n=1). Further, CNE8 and CNE55 (6.3 min, n=1) are both chronic, clade A/E isolates associated with intravenous transmission, thus, we observed no correlation between T20 delay and the diverse clinical characteristics of these isolates.

The clade B transmitter/founder isolates [33] also had typical delays (Fig. 2c) except 62375_14 (2.2 min, n=1), 1006_11 (3.2 ± 1.37 min, SD, n=5) and PRB958_06 (2.4 min, n=1). These early delays were identified among Envs isolated during Fiebig stages II and III, suggesting that early T20 delays may be associated, at a low frequency, with early clinical stages of infection. Like the global panel, several of these isolates have been classified as tier 2 [43] including the typical delay isolates 7000_04, WEAUd15 and SC05, as well as the early delay isolate 1106_11. In addition, the typical delay Env 1012 (6.8 min, n=1) is tier 1B, thus, no clear correlation between T20 delay and reported neutralization sensitivity was observed among these isolates.

To determine whether entry kinetics were associated with transmission, we measured T20 delays for four clade C founder consensus isolates [34, 35] (Fig. 2d). All of these isolates had typical delays ranging from 7.2 to 12.9 min (n=1 for each), suggesting that early founder consensus isolates may exhibit typical delays. Fig. 2e shows isolates associated with mother-to-child transmission [37, 38] (MTCT), where a single isolate was cloned from each infant (Fig. 2e, left panel) and two from each mother (Fig. 2e, right panel). The numbers in each box describe how many Envs isolated from each individual fell into the corresponding T20 delay category. The numbers in parentheses indicate the total number of Envs that were tested from each individual. ML035 harbored a typical isolate (9.2 min, n=1, isolate ML035.G2) and an early isolate that was close to the 4-minute cutoff (3.8 min n=1, isolate ML035.I2). Similarly the other two early isolates were also close to the typical delay threshold with delays of 3.4 min (n=1, isolate BJ613.E1) and 3.8 min (n=1, isolate MI206.D1). All of these isolates are clade A with the exception of the MK184 and BK184 isolates (clade C/D), which all had typical delays, and ML035 (clade D/A). Among these isolates, we did not observe correlations between T20 delay and clade or transmission role. We note that BG505.C2, a prototypical Env for vaccine design, had a typical T20 delay of 8.7 ± 2.9 min (SD, n=4).

The isolates shown so far represent a broad phylogenetic diversity (Fig. 2b), span early clinical stages of HIV-1 infection (Fig. 2c) and are associated with mother-to-child transmission (Fig. 2e). Among the broad clinical, geographic and phylogenetic breadth of these isolates, we observed early delays in 8 of the 44 total Envs (18%). In all cases except the maternal isolates in Fig. 2e, these data represent only a single Env isolated from each individual. Because HIV forms a diverse quasispecies within an individual[30], we wanted to know whether the T20 delays of an individual’s quasispecies was more kinetically diverse.

Fig. 2f shows T20 delay categories for multiple clade C Envs isolated longitudinally from each of five recipients across the first year of infection[36]. The T20 delays of isolates from each individual form an even distribution that resembles the distribution of all isolates combined (Fig 2a). Indeed, the isolates from Z185F, alone, span a range of 30 seconds to 60 minutes. Fig. 2g shows MTCT clade C isolates for which multiple Envs were examined from each individual[31, 39], for which we also observed a broad distribution of T20 delays. Early and late T20 delays were observed among both the maternal and infant Envs, thus, a broad kinetic breadth is not specifically associated with recipient isolates. We do note, however, that extreme late delay isolates were identified in as little as 3 Envs from infants B2010 (38 ± 2.7 min, SD, n=7, B20101_XPR_310, Fig. S3b), B2473 (60 ± 22 min, SD, n=8, B24731_XPD_704, Fig. S3b) and B2661 (50 ± 4 min, SD, n=7, B26611_XPR_217, Fig. S3b). M2661, the paired mother of B2661, also had an extreme late delay isolate of 32 ± 2.6 min (SD, n=4, M26610_PL_114), however, extreme late isolates were not identified in M2010 or M2473. In addition, M1128 harbored one of the earliest delays of 0.5 ± 0.57 min (SD, n=2, M11280_XPR_37, Fig. S2a).

Our results show that the T20 delay spans a >100-fold range, from as early as 30 seconds to 60 minutes. We observed no correlations between T20 delay and clade or Fiebig stage. We found that the T20 delays of isolates representing only a single Env from a single individual were primarily typical (4-6 minutes), however, early delays (2-4 min) were observed at a low frequency (18%). This is consistent with the random chance of selecting an early Env from the broad delay distributions observed among the Envs from Z185F (18%), Z201M (10%) and Z205F (19%). Interestingly, the distribution of delays observed among the 39 Z185F Envs were broader than the 44 combined delays from Fig. 2b to Fig. 2e, suggesting a propensity for obtaining typical or early isolates when only a single Env is isolated. The latest delay observed was an infant isolate from B2473 (60 ± 22 min, SD, n=8), although isolates from B2661 (50 ± 4.2 min, SD, n=7) and Z185F (51 ± 11 min, SD, n=2, Z185F.30JAN03.PL4.1) were similar. Thus, extreme late delays are not restricted to infant Envs. These results help to delineate a broad, natural landscape of entry kinetics where T20 delays range from 30 seconds to 60 minutes and provide a context for understanding the clinical and mechanistic relevance of this diverse phenotype.

### Entry Kinetics Diverge Early After Transmission

The broad T20 delays observed among the clade C heterosexually transmitted isolates (Fig. 2f) and mother-to-child transmitted isolates (Fig. 2g) suggest that a diverse range of kinetics may be associated with the adaptive landscape of an early quasispecies, or with different routes of transmission. Fig. 3a resolves the longitudinal clade C Envs from Z185F, Z153M and Z205F (Fig. 2f) into the approximate month, post infection, from which each isolate was sampled. Z185F had delays of 20.5 ± 0.9 (SD, n=2, Z185F.26OCT02.PB1.1) and 14.6 ± 5 (SD, n=3, Z185F.26OCT02.PB3.1) within the 3rd month post infection and an extreme late isolate was observed at month 6 (51 ± 11 min, SD, n=2, Z185F.30JAN03.PL4.1) (Fig. 3a, top). Z153M harbored a late delay Env of 26 ± 4 min (SD, n=3, Z153M.13MAR02.PL4.1) at the first month of infection (Fig. 3a, middle) and several with delays close to the late range at the third month. Conversely, first-month Z205F isolates included early T20 delays between 2-4 min (Fig. 3a, bottom) and late isolates at months 2 and 7. Although we did not observe a consistent pattern of kinetic adaptation over time, we did find that members of the acute quasispecies can be kinetically diverse.

**Figure 3:**
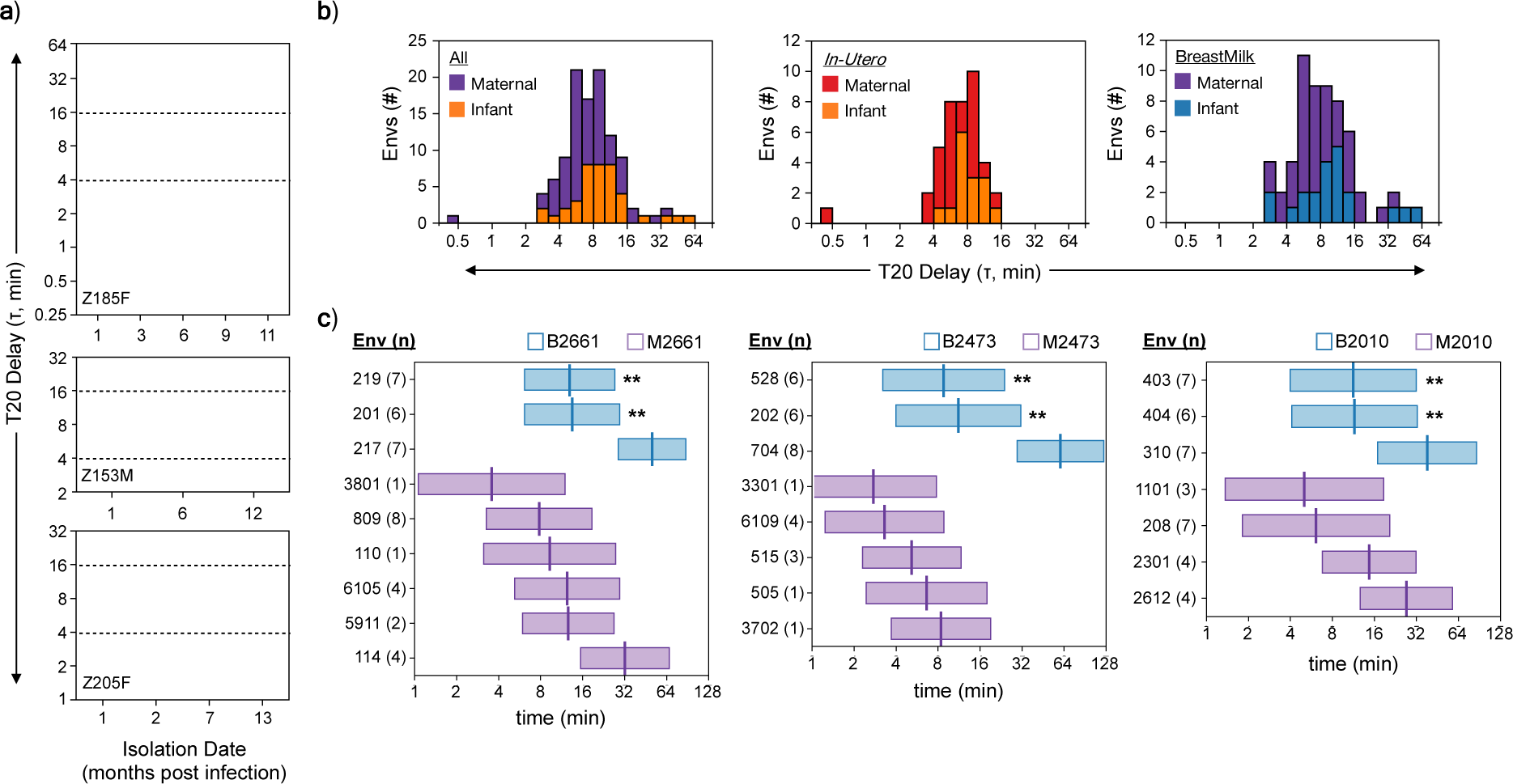
T20 Kinetics Diverge Early After Transmission. (a) T20 delays (τ) for clade C Envs isolated across the first year of infection from individuals Z185F (top), Z153M (middle) and Z205F (bottom). The typical range of 4-16 min is indicated with dashed lines. (b) Stacked histograms of T20 delay for all maternal and infant Zambian isolates (left), isolates associated with in utero transmission (center) or isolates associated with breastmilk transmission (right). (c) T20 kinetic bar profiles for Zambian isolates from mother/infant pairs 2661, 2473 and 2010. The center line of each bar indicates the average delay (τ, min) and the width represents the average duration (D, min). Numbers in parentheses indicate the number of independent experiments and error bars indicate standard deviation where applicable. P values comparing infant Envs to their corresponding extreme late counterpart were calculated using student’s t-test and are indicated as ** p<0.005. All delays shown are the average and standard deviation of 1-8 independent experiments consisting of duplicates, with lognormal R^2^≥0.9.

For the clade C maternal/infant isolates (Fig. 2g) transmission was defined as having occurred either in utero (IUT) or through breastfeeding (BMT) based on the timing of the infant’s first positive PCR test[31, 39]. The transmission route for one mother/infant pair (B2066 and M2066) was indeterminate (IND). While both maternal and infant isolates shared a similar broad range of T20 delay (Fig. 3b, left), the latest delays were infant isolates. When compared by transmission route, the IUT isolates predominantly fell within a narrow 4-16 min window (Fig. 3b, middle), with the exception of the extreme early isolate from M1128 (Fig. S2a). Indeed, the frequency of atypical Envs in the IUT panel (3 in 40, 0.75%) was dramatically lower than the sole frequency of early Envs among the clade C transmission isolates combined (Fig. 2f, 10%) or the isolates from Fig. 2b to Fig. 2e combined (10%). Conversely, the BMT isolates included some of the latest delays (Fig. 3b, right) and spanned nearly the entire range, from early to extreme late.

The latest delays were single isolates from three of six BMT-associated infants. Fig. 3c summarizes the complete T20-refractory transitions using kinetic bar plots that span the average duration of the transition (D, width of each bar, see Fig. 1c) and mark the delay (𝜏, center line in each bar). The delay for B26611_XPR_217 (50 ± 4 min, SD, n=7) was 3-fold later than its relatives, clones 219 (13 ± 3 min, SD, n=7, p=2×10-8) and 201 (14 ± 2 min, SD, n=6, p=2×10-7) (Fig. 3c left). Isolate B24731_XPD_704 (60 ± 22 min, SD, n=8) was 6-fold later than its relatives, 528 (9 ± 3 min, SD, n=6, p=5×10-7) and 202 (11 ± 2 min, SD, n=6, p=6×10-7) (Fig. 3c, center). Isolate B20101_XPR_310 (38 ± 3 min, n=7) was 3-fold later than its relatives, 403 (11 ± 2 min, SD, n=7, p=1×10-6) and 404 (12 ± 5 min, SD, n=6, p=7×10-4) (Fig. 3c, right). Late/extreme late delay isolates from paired mothers were observed for M2661 (clone 114, 32 ± 2.5 min, SD, n=4) and M2010 (clone 2612, 30 ± 16 min, SD, n=4). The remarkably late delays of these infant isolates were confirmed across multiple pseudotype preparations (Fig. S4).

While we did not observe a specific evolution of entry kinetics over time, our data do show a broad range of kinetics among members of acute quasispecies’. Further, we identified one extreme late isolate in only three total isolates that were tested, among 50% of the BMT-associated infants that were examined from this cohort. This is far higher than then 1 extreme late isolate among the 39 Envs from Z185F or the 1 in 6 tested from the paired mother of B2661 (M2661). Conversely, the maternal and infant IUT Envs were much more kinetically confined, with a much lower frequency of atypical isolates than observed overall. These results suggest that entry kinetics can diverge as early as one month post infection and suggest that extreme late isolates may occur more frequently in infants associated with BMT. Previous analyses of the infant Env sequences from this cohort show a high degree of sequence conservation[31], suggesting that these dramatic kinetic differences may be driven by only a few genetic determinants.

### Entry Kinetics Determinants Link Directly to Attachment and Fusion

To identify potential genetic determinants of entry kinetics we compared Env amino acid sequences and T20 delays from the Zambian BMT infants B2661, B2473 and B2010 (Fig. 4). Residues that were unique to late-delay isolates within an infant’s quasispecies were scored as primary candidates (filled arrows) while suspected residues that were not wholly distinguishing were scored as secondary candidates (hollow arrows). B26611_XPR_217 was uniquely distinguished from other B2661 clones by a single substitution A58V (Fig. 4b). All three B2661 isolates with measured delays differed by only 2 additional mismatches: H352Y and N762S (Fig. 4b), which were scored as secondary candidates. B24731_XPD_704 was uniquely distinguished by V570A and R729G, with additional candidates N392S and Q849R (Fig. 4c). The B2010 sequences were more diverse, suggesting multiple determinants. Accordingly, B20101_XPR_310 was uniquely distinguished by three substitutions R327G, I455T and I688V (Fig. 4c) with additional, candidates E351G and G406_01D (an insert near 406).

**Figure 4:**
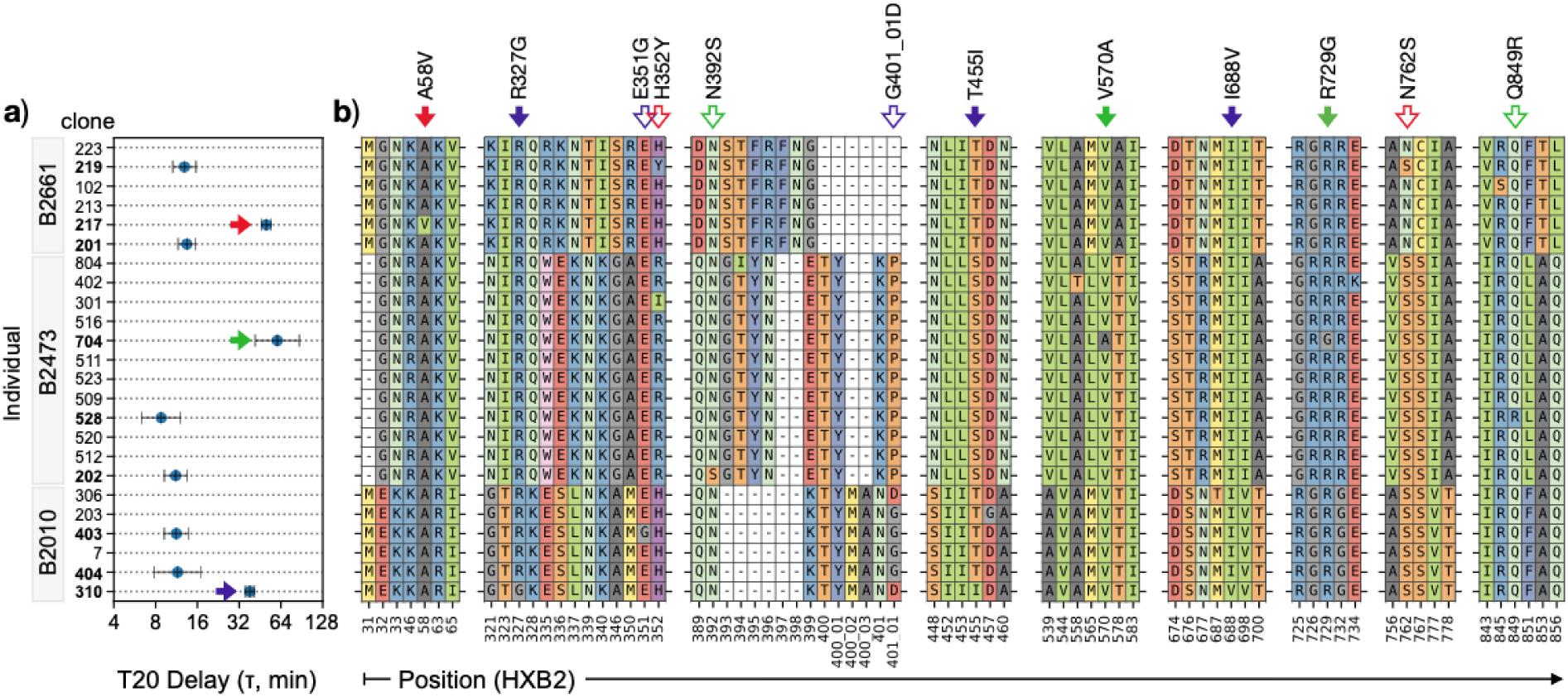
Identification of Kinetic Sequence Determinants. T20 delay, where available (a) and amino acid sequences (b) for Envs isolated from B2661, B2473 and B2010. Arrows indicate extreme, slow isolates for B2661 (red), B2473 (green) and B2010 (violet). Potential sequence determinants of extreme late T20 delays are marked by arrows in panel (b). Solid arrows indicate primary candidates as exclusive distinguishing features of extreme late isolates B2661 (red), B24731 (green) and B2010 (violet). Hollow arrows indicate secondary candidates as differences between related sequences from B2661 (red), B2473 (green) and B2010 (violet) isolates. Amino acid positions follow HXB2 nomenclature. Insert positions are indicated with underscores.

These substitutions are summarized in Table 1, where primary candidates were exclusive to the corresponding late Env and secondary candidates are not nonexclusive, but possible determinants. Using the AnalyzeAlign tool, we found that all primary candidates had low frequency among global HIV sequences (Table 1). To evaluate these candidates we substituted residues from the late delay Envs with those of their early delay counterparts, thus, kinetic determinants will result in a delay that is earlier than the late-delay Env to which the substitution was made.

**Table 1:** Summary of Potential Determinants of T20 Delay. Source Env indicates the isolate from which the residue was identified. Template Env and Substitution indicates the isolate that was mutated and the mutation that was made. Clone substitutions refer to the substitution of a potential late delay determinant with an early delay amino acid. Prevalence was determined using the AnalyzeAlign tool from the Los Alamos HIV sequence database. KE = Kennedy epitope, CD4BS = CD4 binding site, TM = Transmembrane domain. ± indicates standard deviation.

| Source Env | T20 Delay (min) | Residue (HXB2) | Candidate Type | Prevalence | Protein | Location | Template Env | Substitution |
| --- | --- | --- | --- | --- | --- | --- | --- | --- |
| B2661 clone 217 | 50±4 | V58 | Primary | <0.62% | gp120 | Layer 1 | B2661 clone 217 | V58A |
| B2661 clone 219 | 13±3 | Y352 | Secondary | 0.113 | gp120 | α2 | B2661 clone 217 | H352Y |
| B2661 clone 219 | 13±3 | S762 | Secondary | 0.783 | gp41 | KE | B2661 clone 217 | S762N |
| B2473 clone 202 | 11±2 | S392 | Secondary | 0.025 | gp120 | α4 | B2473 clone 704 | N392S |
| B2473 clone 704 | 64±22 | A570 | Primary | <0.38% | gp41 | NHR | B2473 clone 704 | A570V |
| B2473 clone 704 | 64±22 | G729 | Primary | <1.7% | gp41 | KE | B2473 clone 704 | G729R |
| B2473 clone 528 | 9±3 | R849 | Secondary | <0.84% | gp41 | LLP-1 | B2473 clone 704 | Q849R |
| B2010 clone 310 | 38±3 | G327 | Primary | <1.7% | gp120 | V3 | B2010 clone 310 | G327R |
| B2010 clone 403 | 11±2 | G351 | Secondary | 0.037 | gp120 | α2 | B2010 clone 310 | E351G |
| B2010 clone 310 | 38±3 | D406_01 | Secondary | n.a. | gp120 | V4 | B2010 clone 310 | D406_01G |
| B2010 clone 310 | 38±3 | I455 | Primary | 0.017 | gp120 | CD4BS | B2010 clone 310 | I455T |
| B2010 clone 310 | 38±3 | V688 | Primary | <4.1% | gp41 | TM | B2010 clone 310 | V688I |

For B24731_XPD_704, substituting A570V dramatically decreased T20 delay from 60 ± 22 min (SD, n=8) to 11 ± 2 min (SD, n=6, p=10^-7^, Fig. 5a and Fig. 5b), while N392S, G729R and Q849R had no effect. For B20101_XPR_310 (38 ± 3 min, SD, n=7) dramatic changes were observed with I455T (21 ± 2 min, SD, n=5, p=10^-5^) and G327R (17 ± 3 min, SD, n=4, p=0.003) (Fig. 5c, Fig. 5d and Fig. 5e) but not E351G or D406_01G. A58V was the only amino acid difference between B26611_XPR_217 (50 ± 4 min, SD, n=7) and B26611_XPR_201 (13 ± 2 min, SD, n=6), whose T20 delay was significantly earlier (p=10^-7^, Fig. 5e and Fig. 5f). The H352Y and N762S substitutions had no effect on B26611_XPR_217, even across multiple clone preparations (Fig. 5f). The V688I substitution for B20101_XPR_310 did not yield sufficient titers.

**Figure 5:**
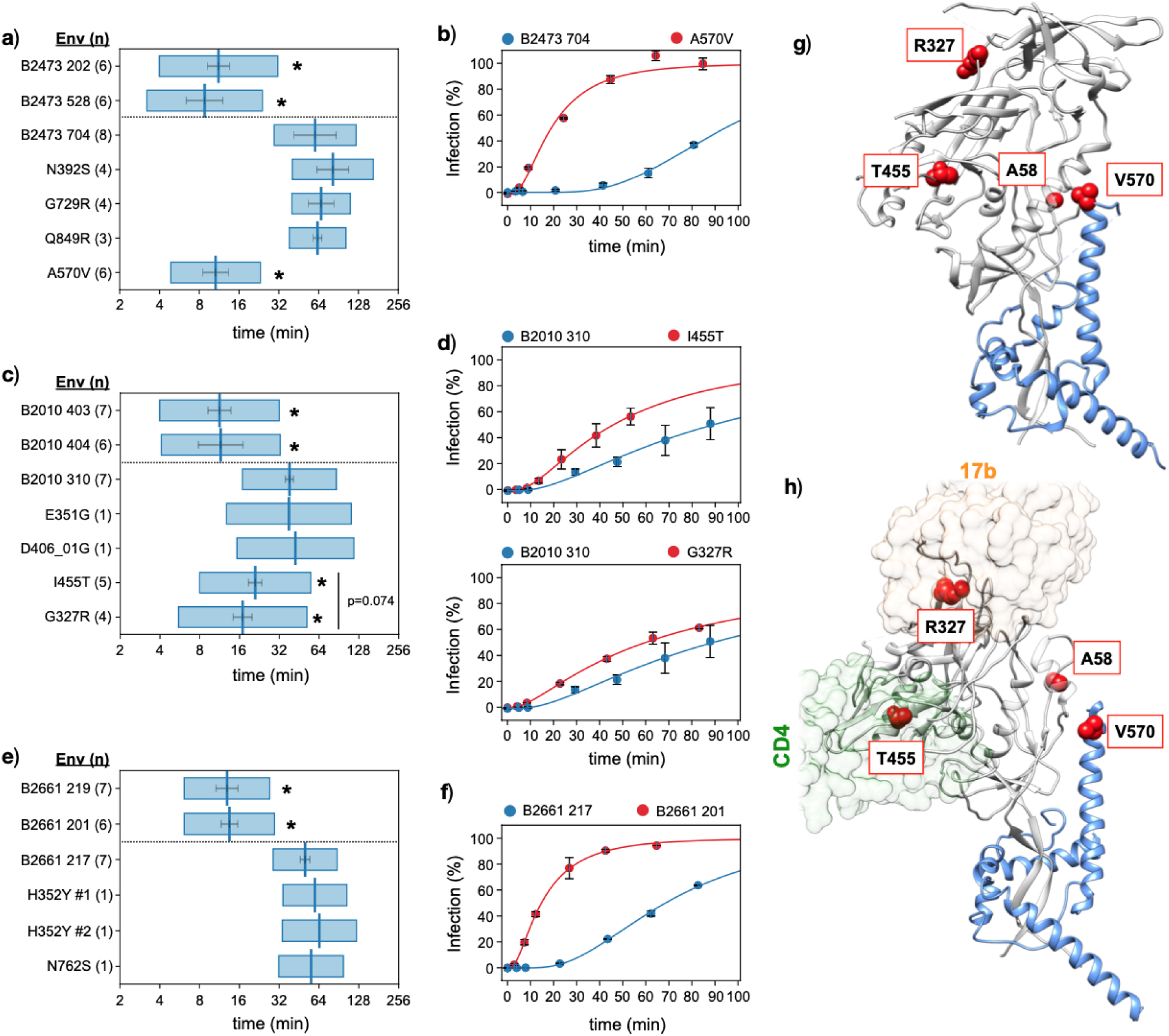
Single Amino Acid Substitutions Dramatically Alter Kinetics. T20 kinetic bar profiles for B2473 (a), B2010 (c) and B2661 (e) isolates and their substituted derivatives. Dashed line divides early delay relatives from the late T20 delay isolate and its substitutions. Representative T20 kinetic curves for B24731_XPD_704 and its A570V substitution (b), B20101_XPR_310 and its I455T and G327R substitutions (d) and B26611_XPR_217 with its V58A relative, clone 201. (g-h) Location of kinetic determinants in (g) the antibody-stabilized pre-fusion trimer structure (PDB ID 6CK9[44]) and (h) the CD4-engaged open trimer structure (PDB ID 5VN3[45]). * isolates with T20 delays that are significantly earlier than the corresponding late delay Env (p<0.0001, calculated with student’s t-test). Numbers in parentheses indicate the number of independent experiments performed for (a, c, e). Circles in (b, d, f) are the average of two duplicates from one representative experiment and error bars are standard deviation. All delays show are from lognormal fits with R^2^≥0.9.

Fig. 5g and Fig. 5h show the positions of significant determinants in both the pre-fusion, closed Env (PDB ID: 6CK9[44]) and the CD4-bound open Env (PDB ID: 5VN3[45]) structures. Residue 58, in layer 1, is linked to anchor inhibitor resistance[46]. Residue 327 is in the third variable loop (V3) and forms a critical CCR5 binding motif [47, 48] that is also a defining epitope for broadly neutralizing V3 antibodies[48–50]. Residue 455 is near the CD4 binding site and affects the affinity of CD4 binding site bnAbs [51] and CD4-Fc[52]. Residue 570 is also associated with anchor inhibitor resistance[53, 54], is a viable neutralizing antibody target [55, 56] and contributes to a wide range of fusion-associated functions including structural stability[57–59], fusogenicity [60, 61] and bundle formation[25, 62, 63]. Overall. These single-residue kinetic determinants fall into two broad functional categories. Residues 327 and 455 are linked to attachment (CD4 and CCR5 engagement), while residues 58 and 570 are linked to gp41-mediated fusion.

### Interdependence Between Co-receptor Engagement and Bundle Formation

The duration of pre-hairpin exposure is an important parameter of fusion inhibitor potency and resistance[19, 20, 46, 64, 65]. While CD4 engagement is sufficient to trigger pre-hairpin exposure[7–11], there is evidence to suggest that co-receptor engagement drives eclipse of the pre-hairpin either through its collapse into the six-helix bundle or through endocytic uptake of the fusing virion [66]. This supported by evidence that low co-receptor density delays entry kinetics and increases sensivity to T20 [19, 20] and that co-receptor antagonists increase sensitivity to fusion inhibitors[22, 67]. To investigate this relationship on a broad scale, we measured delays using maraviroc (MVC), which inhibits co-receptor engagement, for 68 randomly chosen Envs that encompassed a broad range of T20 delay.

Almost half of these MVC delays fell below or near the 2 minute limit of quantitation and were highly variable across experiments (data not shown). Therefore, We focused on 35 Envs whose MVC and T20 delays both had 90% confidence intervals within a 2-fold range. The T20 and MVC delays of these isolates covered similar ranges, from 1 to 60 minutes and were directly proportionate (Fig. 6a, R^2^=8186), suggesting that the MVC delay and T20 delay reflect the same transition.

**Figure 6:**
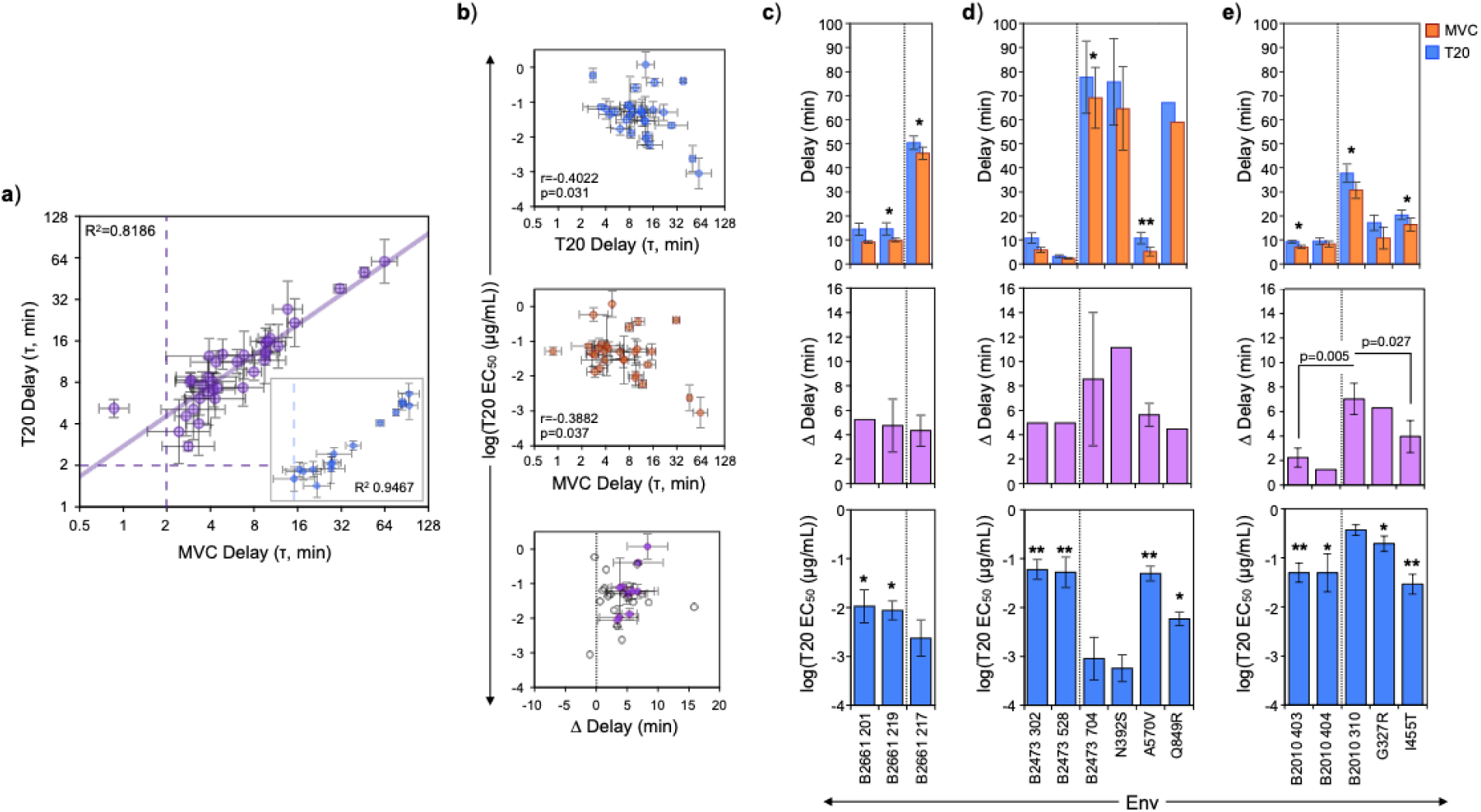
Interdependence Between Co-receptor Engagement and Bundle Formation. (a) Correlation between MVC and T20 delay for 68 diverse HIV-1 isolates (purple circles) excluding the kinetic substitutions. Inset shows MVC and T20 delays for B2661, B2473 and B2010 isolates and their kinetic substitutions. Limits of quantitation are indicated with dashed lines. (b) Correlations of T20 EC_50_ to T20 delay (top), MVC delay (middle) and Δ delay (bottom, T20 delay minus MVC delay). For Δ delay, Envs whose MVC and T20 delays were significantly different (p<0.05) are shown as closed circles with error bars (standard deviation) and those that were not significant are shown as open circles with no error bars. (c-e) paired MVC and T20 delays (top), corresponding Δ delays (middle) and T20 EC_50_ (bottom) for (c) B2661, (d) B2473 and (e) B2010 isolates and their kinetic substitutions. Dashed lines separate early Envs from their late counterpart and its substitutions. Paired MVC and T20 delays are the average of 3-5 simultaneous MVC/T20 experiments. Δ delay are the average and standard deviation of T20-MVC differences. T20 EC_50_s are the average and standard deviation of 3-8 independent experiments involving duplicate with lognormal R^2^≥0.9. Stars for (c-e) (top) indicate significant differences in MVC and T20 delay using student’s t-test. Stars for (c-e) (middle and bottom) are student’s t-test comparisons to the respective, extreme late Envs B26611_XPR_217, B24731_XPD_704 or B20101_XPR_310. All stars indicate p values of * p<0.05, ** p<0.005.

To determine the impact of these kinetics on pre-hairpin exposure, we measured T20 sensitivity for a random subset of 29 of these Envs. Consistent with other results[19–21], T20 EC_50_s formed a weak but significant correlation to T20 delay (Fig. 6b, top, p=0.031). We also observed an equally significant correlation between T20 EC_50_and MVC delay (Fig. 6b, middle, p=0.037). Thus, both T20 and MVC delay are roughly equivalent markers of the duration of pre-hairpin exposure. Despite the similarities between MVC and T20 delay, we did find that MVC delay was 0 to 15 min earlier than T20 delay and this difference was not associated with T20 EC_50_ (Fig. 6b, bottom). This is consistent with a model of entry where pre-hairpin exposure is triggered by CD4 engagement and then eclipsed by co-receptor engagement [66].

The results presented in Fig. 6b support results obtained from a more complete model of entry that uses individual rate constants for each major stage of entry[21]. The complete model demonstrates that T20 sensitivity is proportionate to the average duration of time an Env spends in the CD4-bound and the CD4-bound plus the co-receptor-bound states. These are equivalent to the timing of the MVC and T20 refractory transitions as measured by the lognormal delay, respectively. Similar to this study, we also found that t_50_ (using MVC or T20) did not correlate to T20 sensitivity (Fig. S5).

To examine this relationship in finer detail, we focused on the B2661, B2473, B2010 isolates and their substitutions, which generally belonged to one of two broad categories: positions 570 (gp41) and 58 (gp120) are linked to fusion [46, 57, 58, 60, 61] and positions 327 and 455 are linked to receptor/co-receptor attachment[48, 51, 52]. We performed side-by-side, simultaneous time-of-addition experiments using both MVC and T20 to allow direct and paired comparisons of both refractory transitions (Fig. 6c).

Overall, we observed a strong proportionality between MVC and T20 delay for these substitutions and their wild type counterparts (Fig. 6a inset, R^2^=0.9467). Surprisingly, all of these substitutions had the same effect on both the T20 and MVC delay, regardless of their structural or functional associations (Fig. 6c-e, top). For several of these isolates we were able to statistically distinguish MVC and T20 delays using paired t-tests. Corresponding Δ delays were calculated as the average of individual Δ delays (Fig. 6c-e, middle). These data were combined with T20 EC50 measurements (Fig. 6c-e, bottom) to examine the relationship between co-receptor engagement and pre-hairpin exposure.

Both the T20 and MVC delays of B26611_XPR_217 were roughly 40 minutes later than its closest relatives due to the single A58V substitution in layer 1 (Fig. 6c, top). B26611_XPR_217 had the same Δ delay as its relatives (Fig. 6c, middle) but was 3-fold more sensitive to T20 (Fig. 6c, bottom, p<0.05). This 3-fold increase in sensitivity, associated with a 40 minute increase in MVC and T20 delay is proportionate an extended 30 minute exposure of C34 to temperature arrested X4 and R5 isolates[13].

The T20 and MVC delays for B24731_XPD_704 were approximately 60 minutes later than its closest relatives (Fig. 6d, top). The later delays of B24731_XPD_704 and the N392S substitution were associated with a longer Δ delay (Fig. 6d, middle) and were both 50-fold more sensitive to T20 (Fig. 6d, bottom). The MVC delay, T20 delay, Δ delay and T20 sensitivity of A570V matched the early delay relatives (clones 302 and 528). Interestingly, although the Q849R substitution in LLP1 had little effect on the overall kinetics of B24731_XPD_704, it did result in a shorter Δ delay and a significant 5-fold decrease in T20 sensitivity (Fig. 6d, middle and bottom).

The extreme late isolate B20101_XPR_310 differed from its early delay relatives by two substitutions: I455 falls near the CD4 binding site and G327 is part of the CCR5 binding site. Similar to the B2473 isolates, B20101_XPR_310 had a larger Δ delay than its closest relatives (clone 403 and 404), however, it was 6-fold less sensitive to T20 (Fig. 6e, p<0.05). The I455T substitution significantly reduced MVC delay, T20 delay and Δ delay, and matched the T20 sensitivity of clones 403 and 404. G327R resulted in a partial but significant increase in T20 sensitivity without affecting Δ delay.

Our results show that MVC and T20 delay are proportionate and are both equivalent predictors of T20 sensitivity, suggesting that these delays reflect the same transition. However, there is a discernible temporal difference separating MVC and T20 delays that is not related to pre-hairpin exposure. This suggests that pre-hairpin eclipse is preceded by a brief phase of irreversible co-receptor engagement, suggesting that co-receptor engagement drives pre-hairpin eclipse. We also found that kinetic determinants in gp41 and layer 1, which are not directly linked to co-receptor engagement, can affect both MVC delay and T20 sensitivity. Thus, our data support a mechanism where eclipse of the pre-hairpin is dependent on both co-receptor engagement and fusogenic activity in gp41. While most of our data support a proportionality between T20 sensitivity and slow kinetics, the B2010 isolates oppose this trend, where later delays were associated with T20 resistance.

## Discussion

We used the lognormal delay as a simple distributive model for fitting time of addition kinetics data that is able to recapitulate important relationships connecting T20 sensitivity to MVC and T20 delay. Our careful characterization of HIV entry kinetics from multiple panels of diverse, naturally occurring HIV isolates reveals a broad range of T20 and MVC delays spanning from as early as 30 seconds to as late as 60 minutes. We also identified four single-residue substitutions associated with CD4 engagement, co-receptor engagement and fusogenicity that are, alone, sufficient to confer extreme late T20 and MVC delays. These single substitutions were also linked to significant differences in T20 sensitivity. Finally, careful comparison of MVC and T20 delays with T20 sensitivity reveal an important mutual and functional relationship between co-receptor engagement and fusogenicity.

### The Lognormal Distribution as a Simplified Kinetic Model

The time-of-addition assay is robust in that it is amenable to detailed kinetic modeling, as well as simple distributive modeling to obtain basic timing metrics. The t_50_ metric is convenient because the kinetics of prerequisite states are not needed and because t_50_ can be directly interpolated. We found that the lognormal delay was qualitatively similar to more complex models that have demonstrated a proportionality between T20 sensitivity, co-receptor engagement residence time and pre-hairpin eclipse [21] (as measured by MVC and T20 delay here, Fig. 6b-d). Indeed, we observed this proportionality among a diverse panel of isolates, suggesting a weak, but broadly significant kinetic component to T20 sensitivity that is likely affected by other factors such as binding affinity. We found the lognormal distribution to be a convenient model of intermediate complexity that (a) could be validated against complete kinetic curves, (b) allowed us to examine discrete refractory transitions independently and (c) qualitatively matched more complex and detailed kinetic models.

### Natural Breadth of Entry Kinetics

While some studies have shown relatively homogeneous kinetics [26] among transmitted clade B isolates, we report a very broad range for clade C isolates associated with both heterosexual and mother-to-child transmission. Our results show that individuals can harbor a kinetically diverse quasispecies and that this kinetic breadth can be established in as early as one month post infection. This is supported by our identification of four point mutations that are each significant determinants of entry kinetics. Thus, the kinetic phenotype may have a relatively labile adaptive landscape that might not be reflected in a founder consensus sequence (Fig. 2d). It will be important to know whether the substitutions described here have the same effect in other Envs or whether these effects are context dependent.

The breadth of this phenotype was only apparent when multiple isolates were examined from each individual. This suggests that understanding the biological basis of entry kinetics will require comparisons of the kinetic distributions of multiple members of a quasispecies. For example, we identified one extreme late isolate among a total of three isolates tested, for 50% of the BMT infants in the clade C MTCT cohort (Fig. 2g and Fig. 3c). Conversely, the extreme late Z185F isolate was only 1 out of 39 Envs and the extreme late M2661 isolate was 1 in 6. This suggests that infants associated with BMT may have a higher likelihood of harboring extreme late variants than adults. Prior characterization of these BMT-associated infant Envs showed that they were more sensitive to the V2-glycan broadly neutralizing antibodies PG16 and PG9 and required higher CD4 surface densities to mediate entry[31]. This phenotype suggests an overall high degree of structural stability that could result in a greater frequency of slow kinetic variants than found in adults.

Although a broad range of kinetics was associated with both maternal and infant BMT-associated Envs, the Envs associated with transmission *in utero* had a very narrow kinetic distribution (Fig. 2b). Out of the 40 total IUT isolates only 2 early delays were found. This is a much lower frequency (5%) than was observed among the 44 globally and clinically diverse single isolates (18%) or the kinetically diverse adult transmission isolates (Fig. 6f, 11 total early isolates out of 66, 17%). However, the broad kinetic diversity of the BMT isolates and the narrow range of the IUT isolates was reflected among both maternal and infant Envs. Thus, while entry kinetics do not distinguish maternal and infant isolates from one another, they may reflect selective barriers, or the lack thereof, that are unique to these transmission routes[31].

It will be important to understand how these kinetics relate to replicative fitness. We note that B20101_XPR_301 and B26611_XPR_217 were sequenced from cell-associated RNA, suggesting that these Envs belonged to actively replicating virus[31, 39]. Conversely, B23471_XPD_704 was isolated from DNA and may or may not have been actively replicating. Our data do suggest that extreme early and late kinetic isolates are rare among an individual’s quasispecies. Additionally, it will be important to understand the sensitivity of these rare kinetic extremes to antibody neutralization, which is also associated with structural stability[68]. These extreme kinetic may be capable of seeding new lineages of resistant isolates, despite their apparent rarity.

### Interplay of Co-receptor Engagement and Pre-hairpin Eclipse

The T20 delay corresponds to the eclipse of the pre-hairpin structure either by transition into an early-stage, T20-refractory bundle or by uptake into a T20-inaccessible endosome[13, 14]. CD4 engagement, alone, is sufficient to initiate pre-hairpin exposure[7–11], thus, it is likely that co-receptor engagement drives pre-hairpin eclipse.

Co-receptor-mediated eclipse is supported by evidence that low co-receptor density increases T20 t_50_ and increases T20 sensitivity[19, 20], thus, delayed co-receptor engagement results in a longer duration of pre-hairpin exposure. This same mechanism also explains why co-receptor antagonists increase sensitivity to fusion inhibitors [22–25] whose binding affinity is high enough to be affected by exposure kinetics. Accordingly, we identified a naturally occurring substitution in the co-receptor binding site (G327) that increases both MVC and T20 delays. We also found that MVC and T20 delay could be experimentally distinguished (p<0.05) in several cases (Fig. 6d), suggesting that pre-hairpin collapse is preceded by a brief period of irreversibly bound co-receptor. Further, the mutations and isolates described here could be useful for studying HIV-1 entry mechanics via. single-virion imaging [69], where the lifetime of different states might be increased.

The V570A substitution in gp41 has been shown to confer a greater sensitivity to N and C peptitdes targeting the pre-hairpin by destabilizing 6HB formation [58] while residues in layer 1 proximal to A58 are involved in triggering the CD4-bound state of Env [70]. In addition, we find that these residues are also capable of affecting MVC delay and pre-hairpin exposure. Thus, our results suggest a functional overlap between co-receptor engagement and the fusogenic activity of gp41, where both drive fusion cooperatively. Synergy between co-receptor inhibitors and fusion inhibitors may be driven by a compensatory relationship between fusogenicity and co-receptor engagement. That the substitutions reported here are rare and were discovered among primary, T20-naive Envs suggests that they may have arisen through adaptive pathways that may be associated with co-receptor usage efficiency. In a more general context, our results highlight a functional interplay between fusogenicity and co-receptor engagement that is supported by compensatory resistance pathways against improved, 3rd generation fusion inhibitors[5].

It will be important to evaluate the impact of fusogenic substitutions on co-receptor usage efficiency. Understanding how these substitutions affect neutralization sensitivity can also resolve important structural details associated with metastability and pre-triggering[11, 68]. Indeed, the late delay G327 substitution falls in an epitope that distinguishes subclasses of V3-targeting broadly neutralizing antibodies[48–50], thus, neutralization escape may also have kinetic consequences.

Our data show a general trend where late delay corresponds to increased T20 sensitivity. The B2010 isolates, however, showed the opposite trend. This contradiction was confirmed by the B20101_XPR_310 substitutions, which are associated with CD4 and co-receptor binding, where early delays were linked to increased T20 sensitivity (Fig. 6h). Further studies will be needed to understand the mechanism behind this counterintuitive relationship, which may represent an unexpected pathway of resistance against fusion inhibitors.

### Limitations

Delays <2 min were highly variable across experiments. We attribute this to length of the initiation process, where the earliest achievable time point was 2.5 minutes post initiation. Lognormal fits of these data, therefore, did not capture the initial velocity of refractory infection (compare Fig. 1d top to Fig. 1d middle and bottom). We also acknowledge that the apparent kinetic restriction of IUT-associated maternal/infant isolates (Fig. 3b) may prove to be more diverse with the addition of more isolates. The most kinetically diverse isolates reported here are primarily subtype C, which may be linked to reports that clade C isolates exhibit a greater dependence on CCR5 for entry[27]. Conversely, studies of acute transmission clade B isolates showed little kinetic diversity[26]. We also note, however, that the small panel of clade C founder consensus Envs also had typical delays (Fig. 2d). Our results suggest that investigating the kinetics of multiple isolates from individuals harboring non-C subtypes will be needed to determine clade specificity.

### Conclusion

The entry kinetics of naturally occurring HIV-1 isolates are extremely broad and malleable, with several single-amino acid determinants. The early expansion of entry kinetics after transmission suggest that this phenotype may be an integral part of establishing infection. Our results support a model of entry where eclipse of the CD4-triggered pre-hairpin is a sequential process that begins with co-receptor engagement. However, while show that pre-hairpin eclipse can be delayed by substitutions in the co-receptor and CD4 binding sites, we also show that substitutions in gp41 and layer 1 of gp120 can delay co-receptor engagement. These results suggest a co-operativity between attachment and fusion that sheds light on entry inhibitor synergy and novel resistance pathways.

## Materials and Methods

### Cells and Reagents

T20 and maraviroc (MVC) were obtained from the NIH AIDS Reagent Program, Division of AIDS, NIAID (Cat. 12732 and 11580, respectively) and were prepared according to manufacturers instructions. 293T cells were obtained from ATCC. TZM-bl cells were obtained from the NIH AIDS Reagent Program, Division of AIDS, NIAID (Cat. 8129), courtesy of Dr. John C. Kappes, Dr. Xiaoyun Wu. All cell lines were cultured in D10, consisting of Dulbecco’s Modified Eagle Medium (DMEM, ThermoFisher, Cat. 11965-118) with 10% fetal bovine serum (FBS, ThermoFisher, Cat. 10437-028) and 1% penicillin/streptomycin (ThermoFisher Cat. 15140-163) at 37°C with 5% CO2.

### HIV Env Isolates and Plasmids

Maternal and infant HIV Env isolate plasmids from the Zambia Exclusive Breastfeeding Study were obtained as described[31, 39]. Longitudinal, acute clade C Env isolate plasmids were generously provided by Cindy Derdeyn and Eric Hunter[34–36]. Env isolate plasmids from the Nairobi Breastfeeding Trial [37] (Cat #11674), the global neutralization panel [32] (Cat #12670) and the clade B acute transmitted/founder panel [33] (Cat #11663) were obtained from the NIH AIDS Reagent Program, Division of AIDS, NIAID, NIH. The HIV reporter vector plasmid NL4-3 ΔENV EGFP was obtained from the NIH AIDS Reagent Program, Division of AIDS, NIAID, NIH from Drs. Haili Zhang, Yan Zhou, and Robert Siliciano (Cat #11100)[71].

### Luciferase Assay

Media was aspirated from infected TZM-bl cells on 96 or 384 well culture plates. Either 20μL (96 well) or 12μL (384 well) Glo Lysis Buffer (Promega, Cat #E2661) was added, swished and incubated at room temperature for 5-6 minutes. 5μL lysate was transferred to black 384 well luminescence plates. Luciferase substrate (Promega, Cat #E1501) was prepared according to manufacturer’s instructions. Luminescence was read in counts/s using a Tecan Spark with injector system using 25μL substrate, 2 seconds of orbital shaking and 4 seconds integration.

### HIV Env Pseudotypes

For each pseudotype, 2.5×106 293T cells were seeded onto 10cm tissue culture plates (VWR, Cat #10861-680) in 10mL D10 and incubated for 48 hours. Transfection was carried out using BioT (BioLand Scientific, Cat #B01-01) and a 1:1 molar mixture of the respective Env plasmid with NL4-3 ΔEnv EGFP, following the manufacturer’s instructions. Cells were incubated for an additional 48 hours. Supernatants were harvested and cell debris pelleted at 300g for 5 minutes before decanting, aliquoting 1.1mL into Sarstedt tubes (Fisher Scientific, Cat #50-809-245) and storing at −80°C. Pseudotypes were titered on 384 well plates seeded with 8,000 TZM-bl cells/well in D10 and incubated for 48 hours before measuring luciferase activity.

Time of Inhibitor Addition Assay

We optimized the standard time of inhibitor addition protocol [19, 20, 26–29] to improve cell viability, thermal stability and throughput. Our complete optimized protocol is described in Supplementary Methods.

### Lognormal Fitting

Lognormal cumulative distribution fitting was performed using python 3.6.5 with numpy (1.17.2), pandas (0.24.0) and scipy (1.1.0) to obtain μ and σ parameters, from which delay (τ, Equation 1) and duration (D, Equation 2) were derived as the mode and width (at 60% of max), respectively. The complete process, along with additional metrics and fully documented source code is described in Supplementary Methods.

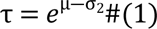

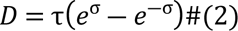

### T20 Sensitivity Assay

TZM-bl cells (8000/well) were seeded onto 384 well tissue culture plates in 50μL D10 and centrifuged for 1 minute at 100g to promote adherence before incubating for 24 hours. T20 stock was thawed and serial dilutions were prepared at four times the final concentration in 12 well reservoirs using D10. 30μL T20 dilutions were aliquoted to 96 well round-bottom plates.

Pseudotype stocks were thawed and diluted to approximately 150,000 to 200,000 RLU, based on TZM-bl titer, final concentration. 90μL diluted virus was aliquoted to the T20 serial dilution plates and mixed. Plates were incubated for 30 minutes at 37°C. For each 96 well plate, 50μL was aliquoted into duplicate wells on a respective 384 well TZM-bl culture plate. Inoculated 384 well plates were incubated for 48 hours before measuring luciferase activity.

### Env Mutation and Sequencing

HIV-1 Env point mutants were prepared using the GeneArt Sited Directed Mutagenesis System (ThermoFisher Cat. A13282). See Supplementary Methods.

### Data Management, Processing and Availability

Experimental data was managed using python 3.6.5 and SQLiteStudio. Calculations and fitting were performed using numpy (1.17.2), pandas (0.25.1) and scipy (1.1.0). The prevalence of potential kinetic determinants was obtained using the Los Alamos HIV sequence database AnalyzeAlign tool https://www.hiv.lanl.gov/content/sequence/ANALYZEALIGN/analyze_align.html. Crystal structure figures were generated using Chimera (1.13.1). Data figures were generated using python/matplotlib (2.2.2), Microsoft Excel (16.15) and arranged using Apple Keynote (9.1).

## Acknowledgements

We would like to thank the authors and participants of all studies whose HIV isolates were used in this study. We thank Drs. Cindy Derdeyn, Eric Hunter and Samantha Burton for providing us with HIV Env plasmids and thoughtful discussions. We thank the NIH AIDS Reagent Program for their outstanding support of our research. We also thank Drs. Otto O. Yang, Tom Chou and Jennifer A. Fulcher for their insights and discussions.

## Author Contributions

Conceptualization: Nicholas E. Webb.

Data Curation: Nicholas E. Webb.

Supervision: Grace M. Aldrovandi and Nicole H. Tobin.

Writing – original draft: Nicholas E. Webb.

Writing -review & editing: Grace M. Aldrovandi and Nicole H. Tobin.

Formal Analysis & Methodology: Nicholas E. Webb.

Investigation: Colin M. Sevareid and Carolina Sanchez.

## Author Declarations

The authors have no conflicts of interest to declare.

